# The regenerating skeletal muscle niche guides muscle stem cell self-renewal

**DOI:** 10.1101/635805

**Authors:** Alicia A. Cutler, Bradley Pawlikowski, Joshua R. Wheeler, Nicole Dalla Betta, Tiffany Elston, Rebecca O’Rourke, Kenneth Jones, Bradley B. Olwin

**Affiliations:** Department of Molecular, Cellular and Developmental Biology, University of Colorado, Boulder, CO, USA; Department of Pediatrics Section of Section of Hematology, Oncology, Bone Marrow Transplant, University of Colorado, Anschutz Medical Campus, Aurora, CO, USA; Departments of Pathology and Neuropathology, Stanford University, Palo Alto, CA

## Abstract

Skeletal muscle stem cells (MuSCs) are essential for muscle regeneration and maintenance. While MuSCs typically are quiescent and reside in an asymmetric niche between the basal lamina and myofiber membrane: to repair or maintain muscle, MuSCs activate, proliferate and differentiate to repair injured tissue, and self-renew to replenish MuSCs. Little is known about the timing of MuSC self-renewal during muscle regeneration and the cellular processes that direct MuSC self-renewal fate decisions. Using DNA-based lineage tracing, we find that during muscle regeneration most MuSCs self-renew from 5-7 days post-injury, following fusion of myogenic cells to regenerate myofibers. Single cell sequencing of the myogenic cells in regenerating muscle reveals that non-cell autonomous signaling networks regulate MuSC self-renewal allowing identification of asymmetrically distributed proteins in self-renewing MuSCs. Cell transplantation experiments verified that the regenerating environment signals MuSC self-renewal. Our results define the critical window for MuSC self-renewal emphasizing the temporal contribution of the regenerative muscle environment on MuSC fate, establishing a new paradigm for restoring the MuSC pool during muscle regeneration.

## Introduction

Skeletal muscle is essential for locomotion, respiration, and longevity and possesses remarkable regenerative capacity dependent on a resident muscle stem cell (MuSC) population (Lepper et al., 2011; Murphy et al., 2011; Sambasivan et al., 2011). MuSCs are quiescent in uninjured muscle and reside in an asymmetric niche bounded on one side by the basal lamina and by the myofiber on the other (Mauro, 1961). In response to muscle injury, MuSCs activate, exit quiescence, and proliferate between 24 and 48 hours post injury (Cornelison & Wold, 1997; Siegel et al., 2011). The resulting myoblasts either differentiate and fuse to regenerate the damaged, multinucleated myofibers or return to quiescence to replenish the MuSC pool.

Replenishing and maintaining a sufficient MuSC pool is critical to sustaining muscle health and regenerative capacity. Muscle regeneration is abolished when MuSCs are lost (Lepper et al., 2011; Murphy et al., 2011; Sambasivan et al., 2011) and decreased MuSC function is associated with aging (Bernet et al., 2014; Collins et al., 2007; Sacco et al., 2010) and myopathies (Heslop et al., 2000). The MuSC population is replenished through symmetric division of a MuSC giving rise to two daughter MuSCs or asymmetric division where one daughter cell inherits proteins associated with activation and proliferation, including MyoD and the other acquires quiescence (Bernet et al., 2014; Conboy & Rando, 2002; Kuang et al., 2007). Signaling pathways influencing MuSC asymmetric division include Notch (Conboy & Rando, 2002; Kuang et al., 2007), Wnt7a (Le Grand et al., 2009), and EGFR (Y. X. Wang et al., 2019) as well as the downstream effectors Vangl2 (Le Grand et al., 2009), p38 (Bernet et al., 2014), and dystrophin (Dumont et al., 2015). These pathways recruit the planar cell polarity pathway to polarize Myf5 and MyoD distribution, thus influencing daughter cell fate. Despite this knowledge, the timing and molecular mechanisms contributing to MuSC self-renewal *in vivo* remain poorly understood as the majority of prior work was performed on cultured myofibers.

Employing *in vivo* DNA base-labeling lineage tracing experiments, we found the majority of MuSCs self-renew between 5 dpi (days post injury) and 10 dpi, following formation of new myofibers, much later than expected from published work examining asymmetric division occurring during the initial cell cycle following MuSC activation in myofiber cultures (Dumont et al., 2015; Kuang et al., 2007; Troy et al., 2012; Y. X. Wang et al., 2019). We performed single cell RNA sequencing (scRNA-Seq) at 4 dpi, the height of myonuclear formation, and at 7 dpi, the height of MuSC self-renewal, which identified coordinated gene expression changes in myogenic cells and self-renewal pathways that collectively drive MuSC self-renewal. We identified major shifts in non-myogenic cell populations in regenerating muscle that may direct MuSC cell fate via non-cell autonomous signaling. During this critical self-renewal window, transplanted MuSC engraftment into the stem cell niche is significantly enhanced, confirming the regenerative environment as a key driver of MuSC self-renewal. The coordinated actions of the regenerating muscle environment prioritize myofiber reconstruction followed by MuSC pool replenishment.

## RESULTS

### The majority of MUSCs self-renew 5-10 days post injury

The timing and kinetics of MuSC expansion, myonuclei generation, and MuSC self-renewal following a muscle injury are largely unexplored. MuSC self-renewal is generally accepted to occur via asymmetric division during the initial cell cycle following MuSC activation (Bernet et al., 2014; Kuang et al., 2007; Troy et al., 2012). If MuSCs self-renew during the initial cell division following quiescence, replenished MuSCs would undergo fewer cell divisions (Fig. 1A), while self-renewal later during regeneration would prioritize myofiber reconstruction (Fig. 1A). To determine when MuSCs self-renew *in vivo*, we lineage-traced MuSCs with timed pulses of 5-Ethynyl-2’-deoxyuridine (EdU) following a muscle injury. If MuSCs self-renew in the initial cell division, we expect lineage labeling of replenished MuSCs from initial EdU pulses. However, if MuSCs self-renew later during regeneration, lineage labeling will not occur with early EdU pulses because initial lineage labeling would be diluted by subsequent cell divisions.

**Figure 1.**
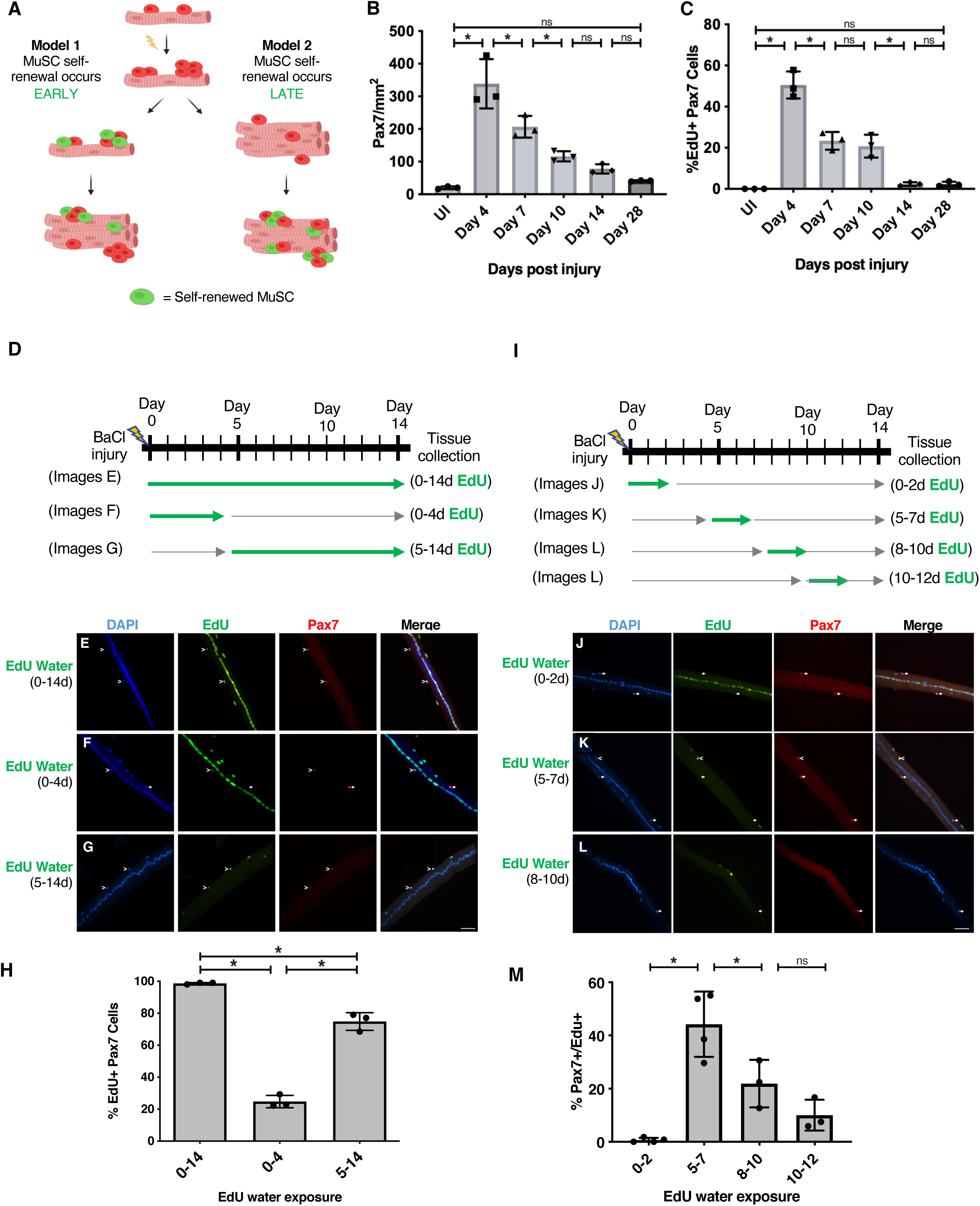
MuSCs primarily self-renew between 5 dpi and 7 dpi. (A) Schematic depicting two simplified models for MuSC self-renewal during regeneration. (B) Pax7+ cell numbers quantified per mm^2^ in tissue sections from injured TA muscle. (C) Pax7+/EdU+ cell percentages in tissue sections from injured TA muscle (D) Experimental design schematic for EdU incorporation DNA-lineage tracing experiments. Green arrows demarcate EdU application timing. (E-G, J-L) Representative images of extensor digitorum longus (EDL) myofibers collected at 14 dpi after the indicated application of EdU. EdU+/Pax7+ double positive cells (white carets) and EdU-/Pax7+ MuSCs (white arrows) are identified. (H) EdU+ MuSCs quantified on myofibers containing central nuclei from exposed to EdU for 0-14 d, 0-4 d, or 5-14 d post injury, respectively (M) EdU+ MuSCs quantified on myofibers containing central nuclei from mice exposed to EdU for 0-2 d, 5-7 d, 8-10 d, or 10-12 d post injury, respectively. All experiments performed as biological triplicates. Statistical significance tested by ANOVA. * denotes p-value < 0.05. All scale bars are 40 μm.

To assess MuSC division kinetics following a muscle injury, we collected tibialis anterior (TA) muscles at various days post BaCl_2_ injury (days 4, 7, 10, 14, and 28). Since DNA base analogs are rapidly diluted (Matiašová et al., 2014), three injections of EdU two hours apart were given immediately prior to tissue collection. MuSCs were identified by Pax7 immunoreactivity and tissue sections processed for EdU incorporation (Supplemental Information Fig S1). Once activated following injury, Pax7+ cells divide rapidly, increasing over 20-fold by 4(dpi then gradually decline to within 2-fold of the initial MuSC number by 28 dpi (Fig 1B). This indicates that most myogenic progenitors are produced early in regeneration and then decline, most likely by terminal differentiation and fusion to form myofibers. Without injury, we detected no Pax7+/EdU+ cells following the three EdU injections prior to muscle harvest (Figure 1C). The highest percentage of Pax7+/EdU+ cells was detected at 4 dpi (50%), which declined to 23% at 7 dpi, and remained at 21% at 10 dpi (Fig 1C). The lowest percentage of Pax7+/Edu+ cells was detected at 14 dpi (2%) with no further decrease by day 28 (Fig 1C). The dramatic decrease in percentage of dividing myogenic cells between 4 dpi and 7 dpi indicates that myogenic cells are leaving the cell cycle either by terminal differentiation or self-renewal. The minimal percent of myogenic cells dividing at day 14 indicates that myonuclei and self-renewed MuSCs are not being produced.

To determine when MuSCs self-renew, mice were given EdU in drinking water for the entirety of the regeneration (0-14 dpi), for the first 4 dpi (0-4 d), or for 5-14 dpi (5-14 d). In all cases, extensor digitorum longus (EDL) muscles were collected at 14 dpi and assessed for EdU incorporation (Schematic in Fig 1D). EdU+ MuSCs and myonuclei were quantified from individual myofibers with centrally located nuclei. MuSCs were identified by Pax7 immunoreactivity and myonuclei were identified as Pax7 negative nuclei located inside the myofiber. Once EdU is incorporated, additional cell divisions without EdU present, dilute the EdU signal until it becomes undetectable; therefore, EdU+ MuSCs and EdU+ myonuclei exited the cell cycle within a few cell divisions of EdU removal. All myonuclei and all MuSCs in myofibers with central nuclei were EdU+ when EdU is provided for 0-14 dpi (Fig 1D, H). In contrast, when EdU is administered from 0-4 dpi, a minority of MuSCs (mean 24%) were EdU+, while all centrally located myonuclei and some peripheral myonuclei were EdU+ (Fig 1F, H). The majority of MuSCs were EdU+ when EdU is administered from 5-14 dpi and while peripheral myonuclei were EdU+, no centrally located myonuclei were EdU+, (Fig 1G, H). We conclude that the majority of MuSCs undergo their final cell division before self-renewing between 5-14 dpi, while the initial MuSC cell divisions predominately produce myonuclei.

To further refine the timeframe of MuSC self-renewal following muscle injury, we administered EdU for shorter durations. We provided EdU water in two-day pulses 0-2, 5-7, 8-10 and 10-12 dpi, harvesting myofibers from the EDL muscles at 14 dpi (Schematic in Fig 1I). On myofibers exposed to EdU water 0-2 dpi, less than 1% of MuSCs were EdU+ and centrally located myonuclei were variably labeled (Fig 1J, M). Since the initial MuSC cell division occurs between 24 and 48 h (Jones et al., 2005; Troy et al., 2012; Webster et al., 2016), few if any MuSCs self-renew during the first cell division following injury, strongly countering the proposed early self-renewal model (Fig 1J) and the majority of published data that assumes asymmetric division, and ergo self-renewal, occurs during the initial division following MuSC activation (Dumont et al., 2015; Kuang et al., 2007; Troy et al., 2012; Y. X. Wang et al., 2019). Furthermore, the variable EdU signal in central myonuclei is consistent with rapidly proliferating myogenic progenitors, diluting EdU following EdU removal at 2 dpi, prior to terminally differentiating (Fig 1J). Approximately 75% of MuSCs self-renew between 5 dpi and 12 dpi with 44% self-renewing from 5 dpi to 7 dpi, 22% self-renewing between 8 dpi and 10 dpi, and 6% self-renewing from 10 dpi to 12 dpi, respectively (Fig 1J-L). The EdU lineage tracing results define the temporal kinetics of fate decisions where the majority of MuSCs self-renew between 5 and 7 dpi.

### A malleable myogenic transcriptome directs myogenic self-renewal

The observation that MuSCs do not self-renew during the initial cell division led us to explore the molecular mechanisms contributing to MuSC self-renewal. We performed single cell RNA sequencing at 4 and 7 dpi, timepoints coinciding with the height of myonuclear production and MuSC self-renewal, respectively. A diverse cellular population comprises regenerating muscle, including assorted immunological cells, fibroadipose progenitors (FAP), and myogenic cells at various stages of activation, differentiation, and self-renewal (Figure 2A).

**Figure 2.**
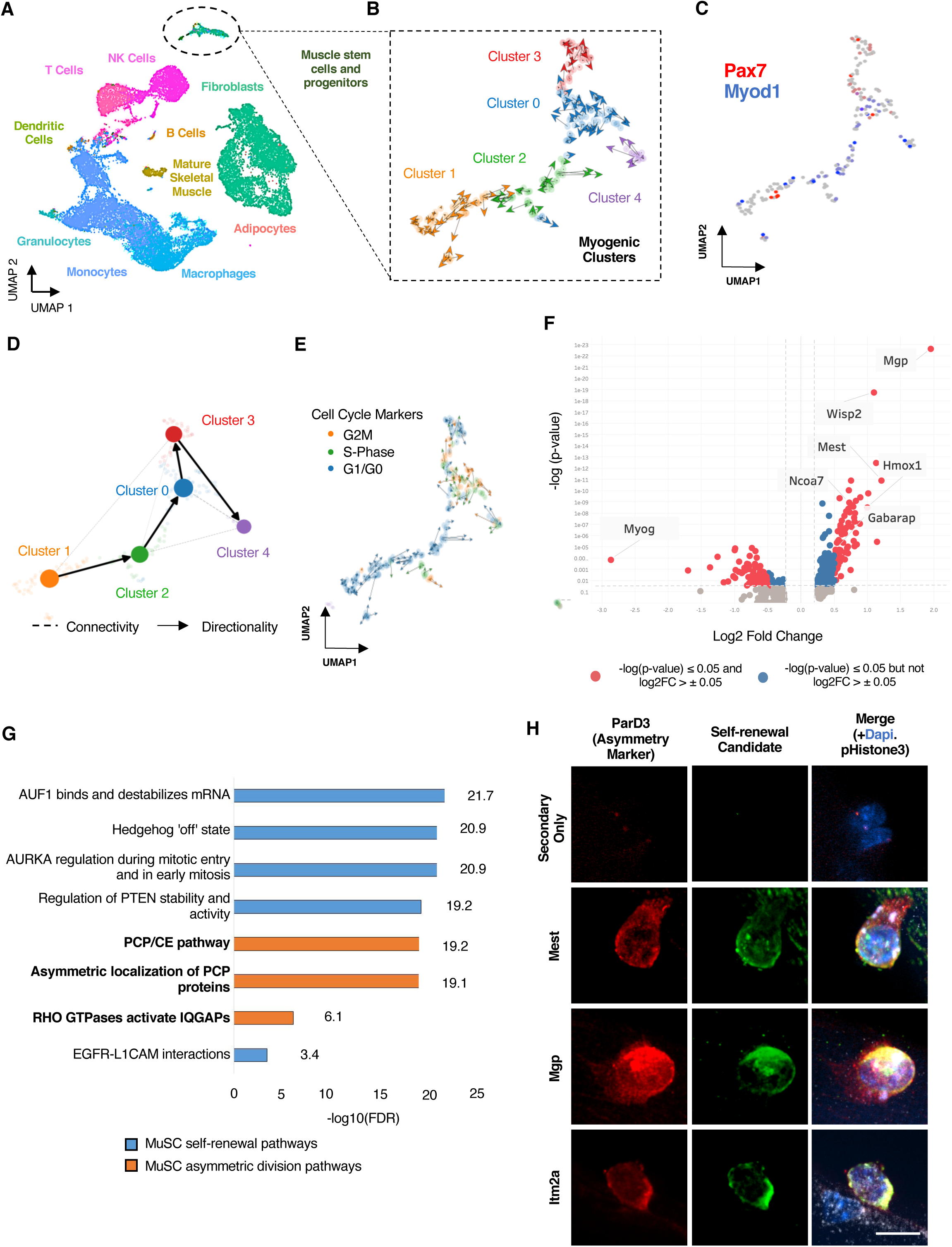
Identifying and characterizing a self-renewing myogenic cluster. (A) UMAP embedding for single cell RNA sequencing of regenerating skeletal muscle at 4 dpi and 7 dpi where clusters are colored by cell type. (B) RNA velocity analysis of myogenic cells at 4 dpi and 7 dpi combined where distinct clusters are indicated by color. (C) UMAP embeddings for Pax7 and Myod1 expression in RNA velocity clusters. (D) Velocity inferred partition-based graph abstraction of myogenic cell fate trajectories. (E) Myogenic clusters from RNA velocity identified by cell cycle stage. (F) Differential gene expression analysis for Cluster 3 (presumed self-renewing myogenic cluster) with candidate genes involved in MuSC self-renewal highlighted. (G) Reactome pathway analysis performed on Cluster 3 differentially expressed genes with self-renewal pathways colored blue and asymmetric division pathways colored orange. Pathways organized by false discovery rate (FDR) (See Table S1). (H) Representative maximum projection images of protein immunoreactivity for ParD3 (red) as a marker of asymmetric division, phosphorylated histone 3 (white) as a marker of dividing cells, and self-renewal candidate proteins (green) as well as DAPI dye (blue) in MuSCs on isolated myofibers.

We focused on defining the molecular features of the mononuclear myogenic population within the regenerative environment. Subclustering the MuSCs and myogenic progenitors identifies five unique MuSC clusters (Figure 2B), each defined by a unique gene signature (Figure S2A). Gene expression analysis revealed cluster-specific expression of genes associated with activation (Pax7 and MyoD) and quiescence (Pax7, Cd34, Myf5) (Figure S2A-B). We identified two clusters (cluster 0 and cluster 3) of Pax7 high, Myod1 low, CD34 expressing MuSCs (Figure 2C, S2B); in cluster 3, Myod1 expression is decreasing and Pax7 expression increasing, suggesting cluster 3 represents self-renewing MuSCs (Figure S2A).

RNA velocity-inferred pseudotime analysis demonstrates a clear directionality of the myogenic clusters towards Cluster 3, the putative self-renewing myogenic cluster (Figure 2D). We initially performed RNA velocity analysis using the scVelo toolkit (Bergen et al., 2020). RNA velocity infers directional trajectories through dynamical modeling of mRNA splicing kinetics. We extended RNA velocity to infer pseudotime of the myogenic clusters by coupling RNA velocity with partition-based graph abstraction (PAGA) to estimate cell-to-cell connectivity while preserving the underlying data topology. We identified a directional hierarchy that is uniformly skewed towards clusters containing proliferating MuSCs and enriched for MuSCs expressing self-renewal and quiescence markers (Figures 2C, S2B). An intermediate population (Cluster 2) that expresses Myod1 RNA (Figures 2C) may represent cycling myogenic cells poised to commit to terminal differentiation or return to quiescence by regulating Myod1 mRNA translation (Hausburg et al., 2015; Morrée et al., 2017). Cluster3, the putative self-renewing cluster, is downstream of the Myod1 high population (cluster 2), further supporting that cluster 3 represents self-renewing MuSCs (Figure 2D).

Finally, MuSCs exit the cell cycle (G1/G0) either to terminally differentiate or to self-renew and a return to quiescence. Of the two G1/G0-positive populations that cell cycle analysis identified (Figure 2E) one correlates with Myod1-positive populations, consistent with these cells exiting the cell cycle and committing to terminal differentiation (Figure 2D, E). The second G1/G0 population is the putative self-renewing cluster further supporting that cluster 3 represents self-renewing MuSCs (Figure 2E).

To interrogate whether cluster 3 contained self-renewing MuSCs and to identify potential drivers of self-renewal we performed differential gene expression analysis on cluster 3 in relation to all myogenic clusters (Figure 2F, Table S1). We further performed differential gene expression analysis comparing cluster 3, the putative self-renewing population with cluster 0, the nearest neighbor of self-renewing MuSCs. We found a collective down-regulation of pro-terminal differentiation myogenic genes and up-regulation of genes potentially involved in self-renewal (Figure S2C, Table S1). Several of the most overrepresented genes from both differential expression analyses are likely related to self-renewal. For example, Mgp, the top result in both differential gene expression analyses (Figures 2F, S2C), influences myostatin expression during muscle development (Ahmad et al., 2017), while Wisp2 (Figure 2F) increases lean muscle mass and muscle Glut4 content (Grünberg et al., 2017). Mest (Figure 2F, S2C) interacts with histone methyltransferases and reducing Mest compromises muscle regeneration following injury (Hiramuki et al., 2015; Zhang et al., 2019). Ncoa7, (Figure 2F) a nuclear coactivator, is required to acidify lysosomes for autophagy (J. Wang et al., 2019) and Gabarap (Figure 2F, S2C) is also involved in autophagy (Fry et al., 2013; H. Wang et al., 2015). Itm2a (Figure S2C), expressed early during skeletal muscle differentiation, promotes autophagy (Lagha et al., 2013; Namkoong et al., 2015). Hmox1 (Figure S2C) is involved in hypoxia response; MuSCs lacking HMOX1 cannot reliably maintain quiescence (Kozakowska et al., 2018). Having defined a potential self-renewal gene expression signature (Ramilowski et al., 2015; Y. Wang et al., 2019), we performed pathway analysis of the differentially expressed genes to identify pathways that may be contributing to self-renewal. Consistent with a myogenic self-renewal program, differential pathway analysis identified proteins like EGFR, AURORA-A kinase, and AUF1, that are components of signaling pathways that regulate MuSC self-renewal in addition to pathways involved in asymmetric cell division (Table S2, Figure 2G) (Chenette et al., 2016; Le Grand et al., 2009; Y. X. Wang et al., 2019).

A subset of MuSC divisions on isolated myofibers are asymmetric with one daughter cell receiving MyoD and continuing to proliferate while the other cell self-renews and acquires quiescence (Bernet et al., 2014; Kuang et al., 2007; Troy et al., 2012). If our scRNA analysis accurately identifies proteins up-regulated in self-renewal, then we expect some these proteins to be asymmetrically partitioned in self-renewing MuSCs. To test this, we selected putative self-renewal target genes from the differentially expressed geneset and validated their expression, subcellular localization, and polarization in MuSCs on isolated myofibers. In asymmetrically dividing cells, identified by phospho-Histone3 (pH3) and asymmetric ParD3, we found six of the seven target genes; Mgp, Wisp2, Hmox1, Ncoa7 and Gabarap; asymmetrically partitioned, while Mest was uniformly distributed (Figure 2H, Supplemental videos). These data define a broader molecular signature for self-renewing MuSCs.

### The influence of regenerating muscle environment on MuSC self-renewal

Most MuSCs self-renew between 5 dpi and 7 dpi when non-myogenic cells and myogenic cells undergo dramatic gene expression changes (Figure 3A, B). We hypothesized that the changing regenerative environment may regulate MuSC cell fate decisions. The composition of the regenerating environment is dynamic and changes in entire cellular populations occurs over three days (Figure 3C). For example, immune cells globally decrease by 15%, while FAPs increase by 10% between day 4 and day 7 (Figure 3D). Consistent with our DNA lineage tracing experiments (Figure 1B), myogenic cells increase between 4 dpi and 7 dpi, indicative of the terminally differentiated progenitors that underwent DNA replication, exited the cell cycle, and are preparing for fusion (Figure 3D). Dynamic fluctuations in the cellular environment may expose muscle progenitors to a pro-differentiation environment or pro-self-renewal environment depending on the signaling presented by cellular interactions and local ligands.

**Figure 3.**
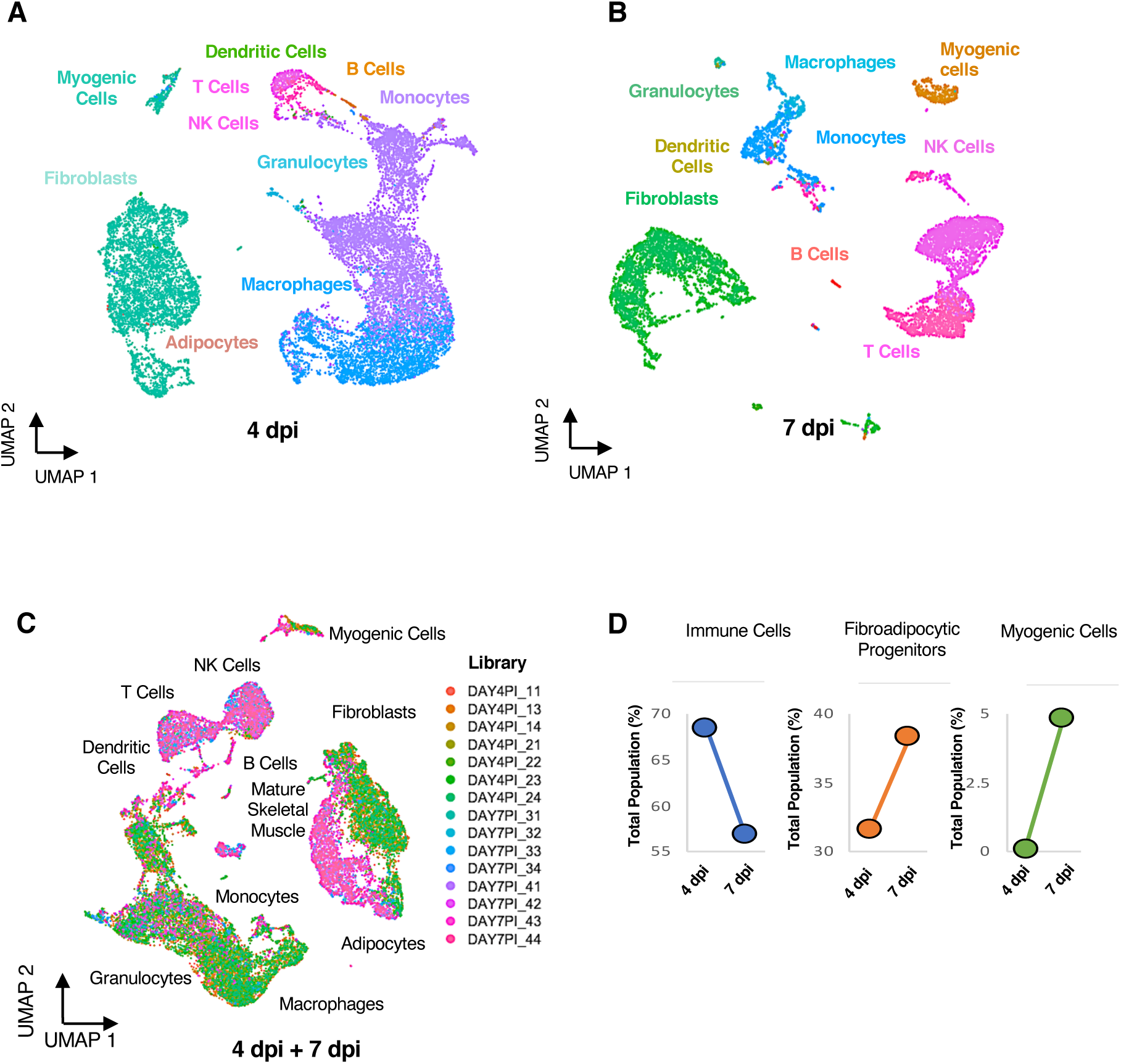
Temporal dynamics of myogenic cells in regenerating muscle. (A) UMAP embeddings of mononuclear cells in regenerating skeletal muscle at 4 dpi and (B) 7 dpi with clusters colored corresponding to cell type. (C) Combined UMAP embedding of mononuclear cells in regenerating skeletal muscle at 4 dpi and 7 dpi with clusters colored by sequencing library. (D) Dynamics changes of mononuclear cells in regenerating skeletal muscle from 4 dpi to 7 dpi for immune cells, FAPs, and MuSCs.

We used NicheNet to map the differential ligand activity during regeneration and examine the impact of these ligand-receptor interactions on muscle progenitors at 4 dpi or 7 dpi (Browaeys et al., 2020). NicheNet draws on *a priori* knowledge of the gene regulatory effects of ligand-receptor interactions on intracellular signaling and gene expression to infer bioactive ligands, identify the cellular source of these ligands, and impute the downstream consequences of ligand-receptor binding on target cell gene expression in an unbiased manner. With stringent cutoffs, we identified differentially expressed ligands predicted to interact with extracellular receptors on muscle progenitors between 4 dpi and 7 dpi (top 20 by differential expression (Figure 4A, Table S3). To determine whether NicheNet accurately mapped cell-MuSC interactions, we queried the NicheNet dataset for described ligand-receptor pairs regulating myogenic cells. Fibronectin 1 (Fn1) is a key adhesion substrate for MuSCs and an essential extracellular matrix adhesion molecule for muscle regeneration (Lukjanenko et al., 2016). NicheNet accurately maps Fibronectin 1 and Fibronectin 1 binding to the Fn1-receptor, B1-integrin (Itgb1) on MuSCs as well as perlecan (Hspg2), meltrin alpha (Adam12) and their respective ligands (Figure 4B, Table S3), all of which are involved in muscle regeneration (Galliano et al., 2000, p. 12; Lafuste et al., 2004; Xu et al., 2010; Yamashita et al., 2018). Thus, NicheNet identifies ligand-receptor interactions and provides predictive power for the temporal impact of the niche on muscle regeneration.

**Figure 4.**
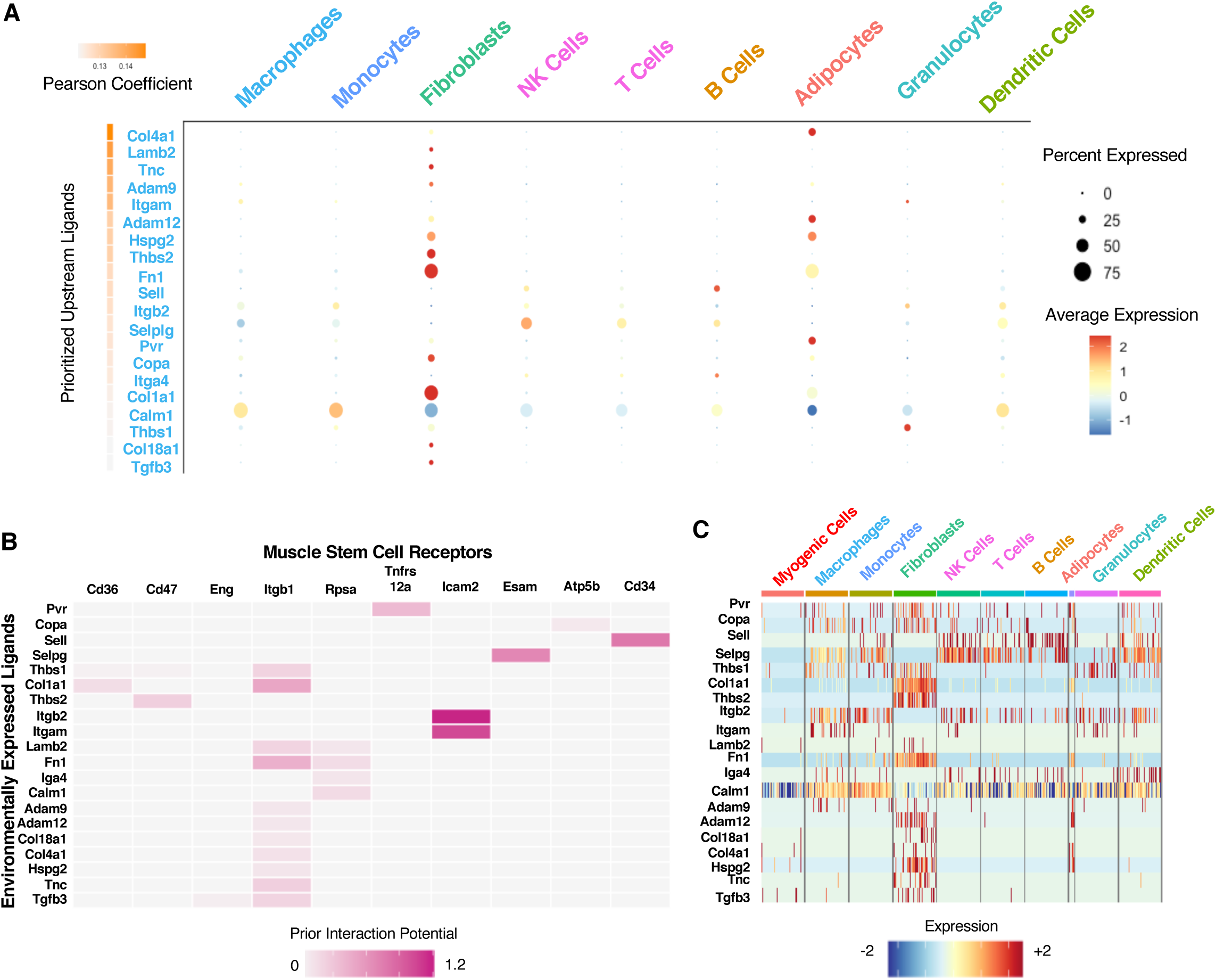
Identifying Signaling Pathways in a self-renewing myogenic cluster. (A) A Dotplot representing NicheNet-imputed ligands produced by mononuclear cells in regenerating skeletal muscle predicted to influence myogenic cells. The dot size reflects the precent gene expression and the hue intensity depicts the relative gene expression. (B) The most significantly expressed genes for cell ligands (by Pearson coefficient) and their potential interactions with genes encoding receptors on MuSCs determined by NicheNet. Hue indicates interaction potential, a representation of the number and quality of data sources supporting the ligand-receptor interaction. (C) Expression levels for the most significant genes encoding ligands in mononuclear cells from regenerating skeletal muscle; the heatmap indicates the relative gene expression level.

Fine mapping of differential ligand-receptor expression during MuSC self-renewal identifies a subset of ligand-receptor interactions that are temporally restricted by ligand expression, receptor expression, or by presence of the ligand producing cells. For example, Thbs1 and Thbs2 (thrombospondin) ligands signal via Cd36 and Cd47, respectively, in myogenic cells (Figure 4B, S4A). Thbs1 is expressed in granulocytes while Thbs2 is predominately expressed in fibroblasts (Figure 4C, S4B). Granulocytes more numerous at 4 dpi, and fibroblasts increase by 7 dpi (Figure 3D). Multiple ligands produced by an array of cells in the regenerating environment may engage the MuSC β1-integrin receptor (Figure 4B, S4A), one example of receptors on myogenic cells capable of binding distinct ligands produced by macrophages, adipocytes, fibroblasts, and dendritic cells (Figure 4C, S4B). Therefore, the changing cellular milieu could provide a complex combinatorial ligand array regulating MuSC fate decisions during muscle regeneration.

### Self-renewal is determined by cell extrinsic factors

To identify additional proteins potentially involved in self-renewal, we examined the expression of receptors identified by NicheNet in myogenic clusters containing self-renewing MuSCs. Hierarchical clustering of molecular signatures for myogenic clusters identifies unique gene expression programs contributing to MuSC self-renewal (Figure 5A). Each myogenic cluster expresses distinct cell surface receptors (Figure 5B); Cluster3 expresses Cd47 and Eng receptors and low levels of Itgb1 and Atp5b receptors. If the environment drives MuSC fates we predict that freshly isolated MuSCs transplanted into a pro-differentiation environment will produce myonuclei but will predominately self-renew when transplanted into a pro-self-renewal environment.

**Figure 5.**
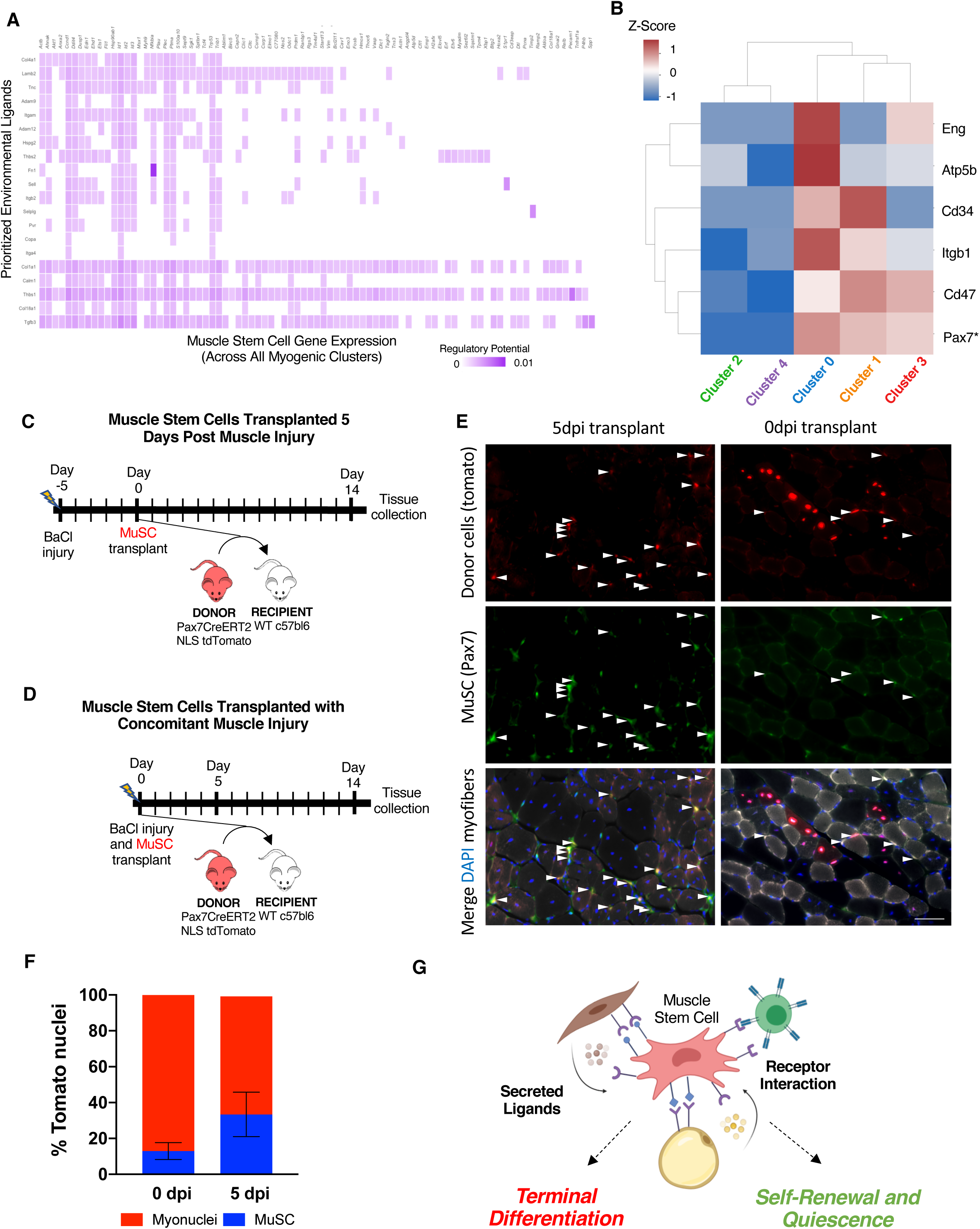
The regenerating muscle environment controls MuSC cell fate. (A) NicheNet imputed regulatory potential of expressed genes encoding ligands for receptors expressed on MuSCs. Regulatory potential represents how well documented the ligand/receptor interactions are and the enrichment of the ligand and receptor transcripts in the dataset. (B) Hierarchical clustering of NicheNet-identified genes encoding receptors on myogenic cell clusters. The heat-map indicates the z-score. (C and D) Schematic of experimental design for MuSC transplantation concomitant with injury, or at 5 dpi (the peak of MuSC self-renewal). (E) Representative images of engrafted, self-renewed tdTomato+/Pax7+ MuSCs (filled arrowheads) at 14 days post-transplant, scale = 100μm (F) The percent of donor cells engrafting either as myonuclei or as MuSCs. Statistical significance tested by t-test with p-value = 0.006 (G) A model depicting the influence of the regenerating muscle microenvironment on MuSC cell fate.

We transplanted freshly isolated MuSCs concomitant with injury or at 5 dpi post injury, the peak of MuSC self-renewal (schematic in Figure 5C, D). MuSCs isolated from mice harboring a nuclear targeted tdTomato transgene preceded by a lox stop lox sequence (Daigle et al., 2018) and a tamoxifen inducible cre recombinase in the Pax7 locus (Murphy et al., 2011) were transplanted into the TA muscles of recipient mice concomitant with injury or at 5 dpi. Muscle tissue was collected at 14 days following transplant and tdTomato positive nuclei quantified as Pax7 positive or negative within the myofiber (Figure 5E). When donor MuSCs were transplanted at 5 dpi, the number of engrafted MuSCs increased 3-fold (Figure 5F) compared to donor MuSCs transplanted concomitant with injury (Figure 5F), which predominately differentiated and fused, becoming myonuclei. Thus, the regenerating muscle environmental appears to direct MuSC fate during skeletal muscle regeneration (Figure 5G).

## DISCUSSION

Replenishing the MuSC pool following injury is essential to maintain long-term muscle health, yet the timing and mechanisms regulating MuSC self-renewal *in vivo* remain largely unknown. Because the MuSC niche is difficult to observe and time consuming to manipulate *in vivo*, the majority of information on MuSC asymmetric division and self-renewal is derived from cultured myofibers (Dumont et al., 2015; Kuang et al., 2007; Troy et al., 2012; Y. X. Wang et al., 2019). We took advantage of DNA based lineage tracing and single cell sequencing to investigate the timing MuSC self-renewal and the mechanisms regulating MuSC self-renewal during muscle regeneration

Lineage tracing during muscle regeneration reveals that most MuSC self-renewal occurs from 5 dpi to 7 dpi, after the majority of myonuclei are produced. In contrast, when EdU is provided from 2 dpi to 4 dpi, myonuclei are labeled nearly exclusively and thus, we identify a critical window when MuSC fate shifts from terminal differentiation to self-renewal. In contrast with data from intact myofiber cultures where MuSC self-renewal is detected during the first cell division following MuSC activation (Dumont et al., 2015; Kuang et al., 2007; Troy et al., 2012; X. Wang et al., 2019), we observe no incorporation of EdU into MuSCs from 0 dpi to 2 dpi, demonstrating MuSC self-renewal does not occur during the first MuSC division post injury. However, we detect heterogeneity for MuSC self-renewal timing where >75% of MuSCs self-renew after 5 dpi, yet a minority of (a mean of 24%), self-renew between 2 dpi and 4 dpi. MuSC heterogeneity in gene expression and degree of quiescence has identified rare populations of reserve MuSCs maintaining quiescence following an injury (Chakkalakal et al., 2012, 2014; Kuang et al., 2007) and identified an alert MuSC state in uninjured muscle (Rodgers et al., 2014). What directs this minority of MuSCs to self-renewal in an environment promoting proliferation and differentiation is unclear. The early self-renewing MuSCs could arise from cell intrinsic differences or environmental differences directing select MuSCs to self-renew.

Detailed, mechanistic examinations of asymmetric division performed by our group and others have focused on cultured myofibers with associated MuSCs for the sake of practical accessibility. However, isolated myofibers lack the basal lamina, half of the MuSC niche, and the milleu of signals in the regenerating muscle environment and thus, there is an unmet need for developing *in vivo* tools to elucidate the mechanisms involved in MuSC self-renewal. We identified self-renewing MuSCs *in vivo* during regeneration by subclustering myogenic cells from data obtained by single cell sequencing at the height of myonuclear production and at the height of MuSC self-renewal. Cluster 3 contains non-dividing Pax7 high/Myod1 low cells and is up-regulated for genes encoding proteins involved in MuSC self-renewal including Aurora Kinase A, EGFR, GSK, JAK signaling (Chenette et al., 2016; Le Grand et al., 2009; Y. X. Wang et al., 2019), Auf1 (Chenette et al., 2016) and Pcp-wnt (Le Grand et al., 2009). By comparing single cell sequencing at the height of myonuclear production and at the height of MuSC self-renewal, we identified signaling pathways that likely regulate these cell fate decisions. Among the ligand-receptor pairs we identified are those with significant overlap with ligand-receptors pairs identified in mice and in human muscle (p = 2.5×10^-18^) using alternative computational methods (De Micheli, Laurilliard, et al., 2020; De Micheli, Spector, et al., 2020). Further, we identified modulators of autophagy and DNA damage repair likely involved in MuSC self-renewal that establish and maintain quiescence (Gassen et al., 2019; Liu et al., 2018; Umansky et al., 2015; Yue et al., 2017). We identified several receptor-ligand pairs yet to be implicated in self-renewal. Among these receptor-ligand pairs are perlecan (Hspg2), Adam12, and Laminin receptor (Lamr/Rpsa). Perlecan, a component of the basal lamina involved in mechanosensing and regulating myostatin expression, muscle size and composition, is upregulated during self-renewal likely modulating MuSC cell fate (Xu et al., 2010; Yamashita et al., 2018). Adam12, a matrix metalloprotease with intriguingly contradictory roles, is upregulated during MuSC differentiation and is required for fusion (Galliano et al., 2000). Surprisingly, over-expression of Adam12 drives cultured cells to adopt a quiescent-like reserve state (Cao et al., 2003) and identify Adam12 as an attractive target to determine its role in MuSC self-renewal *in vivo*. Lamr/Rpsa regulates cell proliferation and migration in non-myogenic cells; as a ribosomal associated protein, Lamar/Rpsa directly links receptor signaling to RNA translation (Scheiman et al., 2010). We identified several up-regulated genes encoding intracellular proteins implicated in stem cell functions of non-myogenic cells. A number of these proteins are polarized in asymmetrically dividing MuSCs identifying potential new regulators of MuSC self-renewal and providingtools to examine MuSC self-renewal *in vivo*.

Identifying genes encoding ligand-receptor pairs that may provide information for partitioning asymmetric distribution of intracellular proteins implicated in stem cell function supports the premise that MuSC self-renewal is driven by the regenerating muscle environment. If the environment is a primary driver for MuSC self-renewal then naïve MuSCs exposed to the self-renewing environment should self-renew, limiting their expansion and production of myonuclei. By transplanting freshly isolated MuSC donor cells into either 5dpi or freshly injured muscle, we confirmed the environment profoundly influences MuSC fate, supporting our premise that environmental signaling drives MuSC fate decisions. When MuSCs are transplanted concomitant with a muscle injury the cells expand as progenitors and primarily engraft as myonuclei. In contrast, when transplanted into the self-renewing environment at 5dpi, the majority of donor cells engraft as MuSCs. These findings have implications for cell-based therapies where a major hurdle is encouraging transplanted cells to engraft as both myonuclei, contributing to healthier, repaired myofibers, and engraft as MuSCs prepared to maintain the muscle for years to come. By manipulating the signals encountered by MuSCs upon transplant, a greater percent of donor cells could be encouraged to adopt a MuSC fate, ensuring longevity of treatment. Moreover, MuSC fate appears more flexible than previously appreciated. By providing different signals in the regenerating environment a single population of freshly isolated MuSCs adopts different fates, either undergoing terminal differentiation or self-renewing.

A dynamic regenerative environment promotes efficient skeletal muscle regeneration by inducing rapid expansion of activated MuSC and initially promoting differentiation of MuSC progeny to produce myonuclei evolving to a pro-self-renewal environment between 5 dpi 7 dpi. We identified ligand receptor pairs, signaling pathways, and intracellular proteins present at these distinct regenerative phases supporting the conclusion that environmental cues direct MuSC fate. The proteins we found up-regulated in self-renewing cells and asymmetrically distributed provide novel tools to further probe the mechanisms responsible for MuSC self-renewal and the signaling pathways that drive cell fate decisions in the regenerative muscle environment.

## Experimental Procedures

### Mice

Mice were bred and housed according to National Institutes of Health (NIH) guidelines for the ethical treatment of animals in a pathogen-free facility at the University of Colorado at Boulder. University of Colorado Institutional Animal Care and Use Committee (IACUC) approved animal protocols and procedures. The mice were C57B6 (Jackson Labs Stock No. 000664) or Pax7-cre^ERT2^ NLS-TdTomato (Jackson Labs Stock No. 025106) between 4 and 8 months old and were a mix of male and female.

### Mouse Injuries and EdU delivery

Mice were anesthetized with isoflurane followed by injection with 501L of 1.2% BaCl_2_ in normal saline into the TA and EDL muscle. To deliver EdU, mice were either given water containing 0.5mg/ml EdU (Carbosynth), with 1% glucose or given IP injections of 10 mM EdU (Carbosynth), re-suspended in water, a volume of 100 µL per 25g mouse weight.

### Myofiber isolation, immunostaining, and culture

The EDL muscles were dissected and placed into 400 U/mL collagenase at 37°C for 1.5h with rotation and then transferred into Ham’s F-12C and 15% horse serum to inactivate the collagenase. Individual EDL myofibers were separated using a glass pipet and immediately fixed using 4% paraformaldehyde for 10 min at room temperature and stored in PBS at 4°C. For visualization of EdU incorporation and immunocytochemistry, myofibers were permeabilized with 0.5% Triton-X100 in PBS (phosphate buffered saline without calcium or magnesium pH 7.4), containing 3% bovine serum albumin (Sigma) for 30 min at room temperature. EdU incorporation was visualized using the Click-iT EdU Alexa Fluor 488 imaging kit (ThermoFisher) following the manufacturer’s protocol. Primary antibodies [anti-Pax7, 2μg/mL (Developmental Studies Hybridoma Bank (DSHB) at the University of Iowa), anti-mgp 1:100 (Proteintech), anti-ncoa7 1:100 (Novus Biologicals), anti-mest 1:100 (Invitrogen), anti-itm2a (Proteintech), anti-HO-1 1:100 (Proteintech), anti-wisp2 1:100 (Invitrogen), anti-gabarap1 1:100 (abcam), anti-parD3 1:200 (Santa Cruz Biotechnology), anti-phospho histone 3 1:500 (Millipore)] were incubated with intact myofibers at room temperature for 1h followed by three washes in PBS. Myofibers were then incubated with appropriate fluorescently conjugated secondary antibodies [Donkey anti-mouse IgG1 (ThermoFisher), anti-rat (ThermoFisher), anti-goat (ThermoFisher), anti-mouse IgG2a (ThermoFisher)] diluted 1:1000 for 1hr at room temperature. Myofibers were washed 3 times in PBS, incubated with 1 μg/mL DAPI for 10 min at room temperature to label nuclei and then mounted in Mowiol supplemented with DABCO (Sigma-Aldrich) as an anti-fade agent.

### TA muscle collections and cell isolations

TA muscles were dissected and placed into 400U/mL collagenase at 37°C for 1h with shaking and then placed into Ham’s F-12C supplemented with 15% horse serum to inactivate the collagenase. Cells were passed through three strainers of 100µm, 70µm, and 40µm (BD Falcon) and flow through was centrifuged at 1500×g for 5 min and the cell pellets were re-suspended in Ham’s F-12C. To remove dead cells and debris, cells were passed over the Miltenyi, dead cell removal kit columns (Cat# 130-090-101). To remove red blood cells (RBC), cells were incubated with antiTer119 micro magnetic beads and passed over a Miltenyi column (Cat#130-049-901). For the adult and aged uninjured TAs, 6 TA muscles (from 3 mice) were pooled together. For the injured TA muscles 2 TA muscles from 2 different mice were pooled together. Cells were then counted using a BioRad TC20 automated cell counter and processed with the 10X genomics single cell sequencing kit.

### Single Cell sequencing

To capture, label, and generate transcriptome libraries of individual cells we used the 10X genomics Chromium Single Cell 3’ Library and Gel Bead Kit v2 (Cat #PN-120237) following the manufactures protocols. Briefly, the single cell suspension, RT PCR master mix, gel beads, and partitioning oil were loaded into a Single Cell A Chip 10 genomics chip, placed into the chromium controller, and the chromium Single Cell A program was run to generate GEMs (Gel Bead-In-EMulsion) that contain RT-PCR enzymes, cell lysates and primers for illumine sequencing, barcoding, and poly-DT sequences. GEMs are then transferred to PCR tubes and the RT-PCR reaction is run to generate barcoded single cell identified cDNA. Barcoded cDNA is used to make sequencing libraries for analysis with Illumina sequencing. We captured 16,868 from the 4-dpi and 8,995 from the 7-dpi. Sequencing was completed on an Illumina NovaSeq 6000, using paired end 150 cycle 2x150 reads by the genomics and microarray core at the University of Colorado Anschutz Medical Campus.

### Single Cell Informatics

CellRanger version 3.0.1 (10X Genomics, Pleasanton, CA) was used for initial processing of sequencing reads in keeping with standardized workflows with reads mapped to the mouse reference transcriptome mm10. For each library, gene expression matrixes generated, and subsequent downstream analysis was conducted using either R version 3.6.2 (2019-12-15) or R version 4.0.0 (2020-04-24). For each library, quality control, filtering, data clustering and visualization was carried out using Seurat version 3.1.5 R package with some custom modifications to the standard pipeline (Stuart et al., 2019). Each dataset was independently analyzed and datasets from the same time points were combined for integrated analysis. Briefly, for each individual dataset, pre-processing was conducted according to standard workflows wherein genes that were expressed in less that 3 cells as well as single cells harboring less than 2500 UMIs, greater than 5% UMIs mapping to mitochondrial genes, or less than 200 genes were removed from the gene expression matrix. Doublets were identified and removed using DoubletFinder (McGinnis et al., 2019). After log-normalizing the data, we performed PCA on the gene expression matrix and used the first 12 principal components for clustering and visualization. Unsupervised shared nearest neighbor (SNN) clustering was performed with a resolution of 0.5 and visualization was done using uniform manifold approximation and projection (UMAP). Cell identities were annotated using SingleR version 1.2.3 (Aran et al., 2019). Differential gene expression analysis was performed using either Seurat version 3.1.5 “FindMarker” or the DESeq2 package. Data was plotted using Seurat enabled ggplot2, custom R ggplot2 scripts, or exported for graphical analysis using Seaborn version 0.11.0 in Python or Tableau visualization software.

### RNA Velocity and Trajectory Mapping

RNA velocity was performed using the scVelo package (Bergen et al., 2020). In brief, using Python version 3.6.3, velocyto.py alongside Samtools version 1.8 was used to generate loom files pertaining to spliced and unspliced information from CellRanger version 3.0.1 generated output files. Loom files for each Seurat object from each processed library were created using LoomR version 0.2.0 and combined with velocyto output loom files into an adata object using custom Python scripts. Subsequent RNA velocity analysis along with velocity-inferred trajectory and connectivity mapping were performed using standardized scripts with the outputs superimposed onto the UMAP embeddings.

### Ligand-Receptor Interaction Mapping

Ligand-Receptor interaction analysis was conducted using NicheNet using the R-enabled nichenetr plugin (Browaeys et al., 2020) and the Seurat version 3.1.5 R package in R version 4.0.0 (2020-04-24). In brief, a combined processed Seurat object was created for 4 and 7-dpi with embedded cell annotations having been generated using SingleR as above. Metadata was amended to include either a 4-dpi or 7-dpi descriptor for each cell. Previously generated weighted network files were read in and genes were converted to mouse orthologs based on one-to-one orthology. NicheNet analysis and differential expression was performed using standard scripts with “receiver” cells defined as myogenic clusters and “sender” cells defined as all remaining SingleR-annotated cell clusters. Data was plotted using Seurat enabled ggplot2 or exported for graphical analysis using Seaborn version 0.11.0 in Python.

### MuSC transplants

For MuSC transplants, *Pax7^CreERT2^; R26R^NLStdTom^* mice were injected with 100ul 20mg/ml tamoxifen once daily for five days to induce recombination and expression of tdTomato in Pax7 cells. These tomato positive donor mice were then sacrificed and MuSCs collected from hindlimb muscles as described above. To remove dead cells and debris, cells were passed over the Miltenyi, dead cell removal kit columns (Cat# 130-090-101). To remove RBCs, cells were incubated with antiTer119 micro magnetic beads and passed over a Miltenyi column (Cat#130-049-901). To deplete non-MuSCs, cells were incubated with satellite cell isolation kit micro magnetic beads and passed over a Miltenyi column (Cat# 130-104-268). The resulting purified MuSCs were separated into 100,000 cell aliquots and resuspended in either normal saline (for 5dpi transplants) or 1.2% BaCl_2_ in normal saline (for 0dpi transplants) and injected into the TA of an anesthetized mouse.

### Microscopy and image analysis

All images were captured on a Nikon inverted spinning disk confocal microscope. Objectives used on the Nikon were: 10x/0.45NA Plan Apo, 20x/0.75NA Plan Apo, and 60x/1.4NA Plan Apo. Images were processed using Fiji ImageJ. Confocal stacks were projected as maximum intensity images for each channel and merged into a single image. Brightness and contrast were adjusted for the entire image as necessary.

### Statistics

Statistical analysis was performed in Prism (GraphPad), where statistical significance was assessed using, two-tailed, unpaired Student’s *t* test or one-way ANOVA with *p* < 0.05 considered significant. Each N was generated from an individual mouse.

## Author Contributions

BP, AAC, and BBO conceived the experiments. BP, AAC, ND, and TA performed the experiments, analyzed data, and made figures. JW, KJ, and RO performed bioinformatic analysis of single cell sequencing. AAC, JW, BP, and BBO wrote the manuscript. BBO supervised the research. All authors read and approved the manuscript.

BBO discloses a potential conflict of interest as a Scientific Advisory Board Member for Satellos Biosciences

## Data and materials availability

All data is available in the main text or the supplementary materials. Sequencing data deposited as follows: 10X scRNA sequencing (GEO accession number PRJNA#*)

---

**Supplemental Figure 1.**
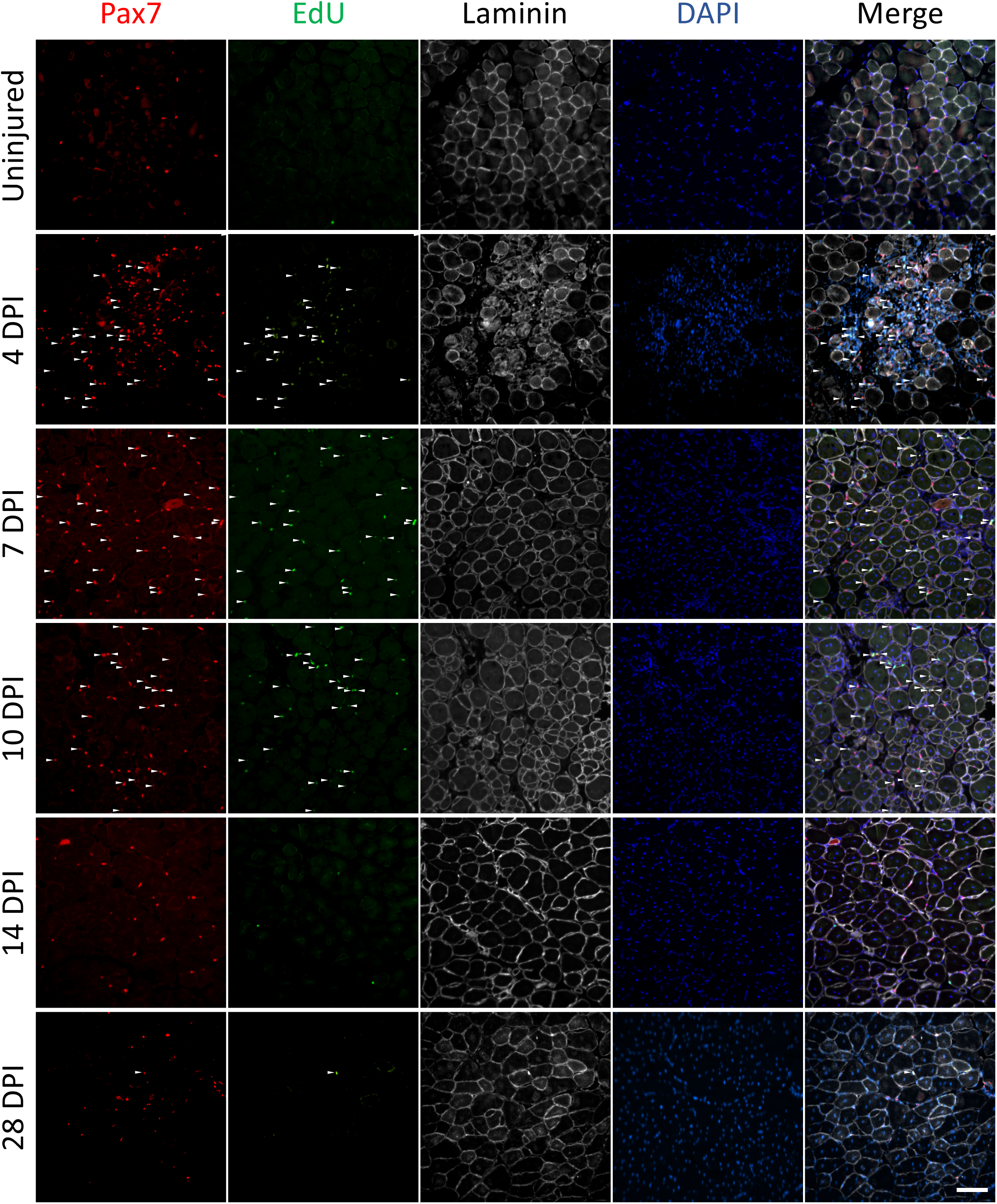
Related to Figure 1. (A) Representative tissue sections from injured TA muscles at the indicated dpi. Three EdU injections two hours apart were given prior to collection. MuSCs (white arrows) are Pax7immunoreactive (red) and EdU+ (green) if labeled by EdU injection., Myofibers are outlined by laminin immunoreactivity (white) and nuclei identified by DAPI staining. All scale bars are 100 μm.

**Supplemental Figure 2.**
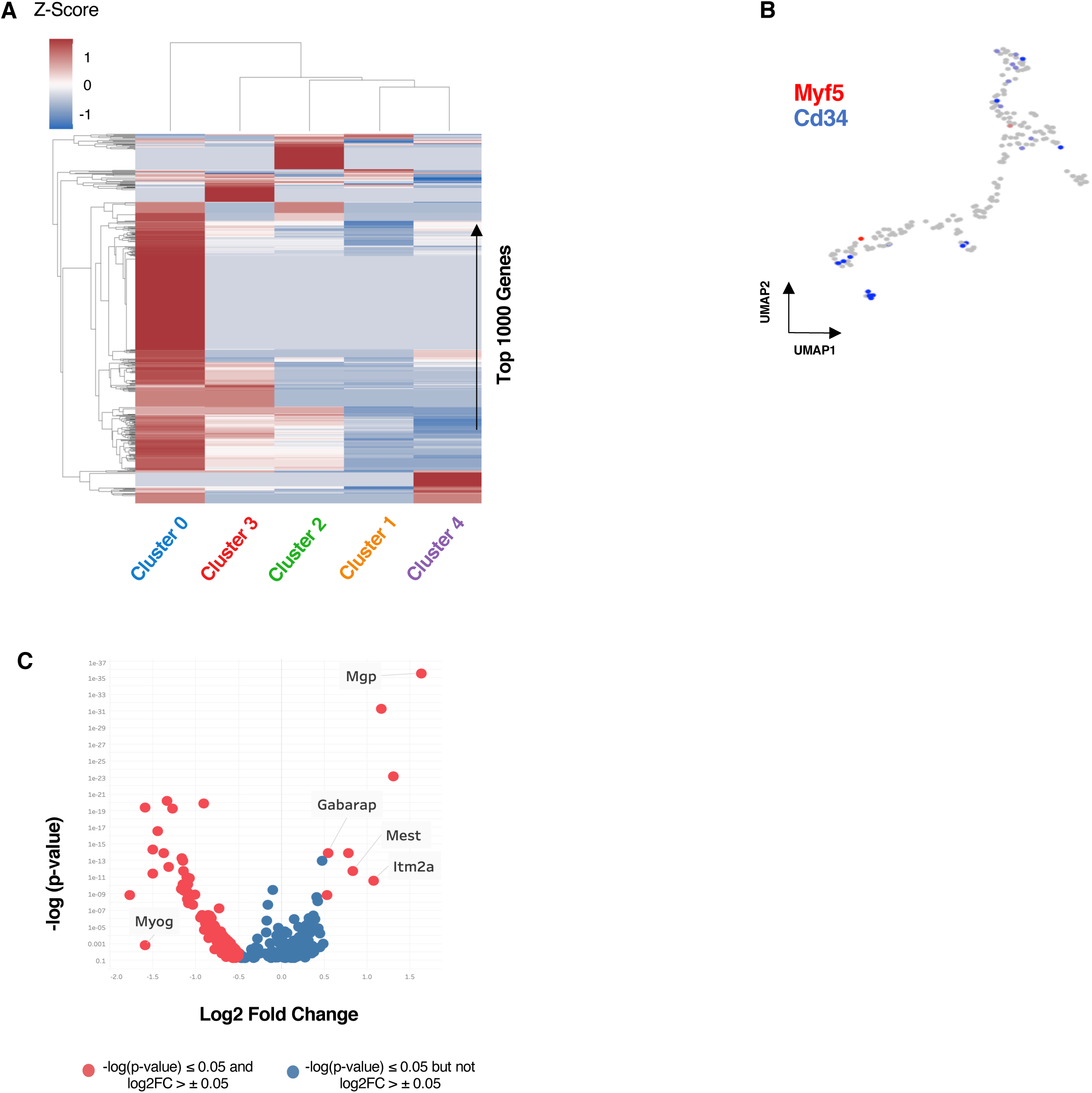
Related to Figure 2. (A) Hierarchical clustering of the 1000 most highly expressed genes across each myogenic cluster. Heat map indicates z-score and relative expression with red indicating increased expression, blue decreased expression, and gray undetected. (B) UMAP embeddings for Myf5 and Cd34 expression in identified myogenic progenitor cell populations. (C) Differential gene expression analysis for Cluster 3 compared to nearest neighbor, Cluster 0, with presumptive genes involved in MuSC self-renewal highlighted.

**Supplemental Figure 3.**
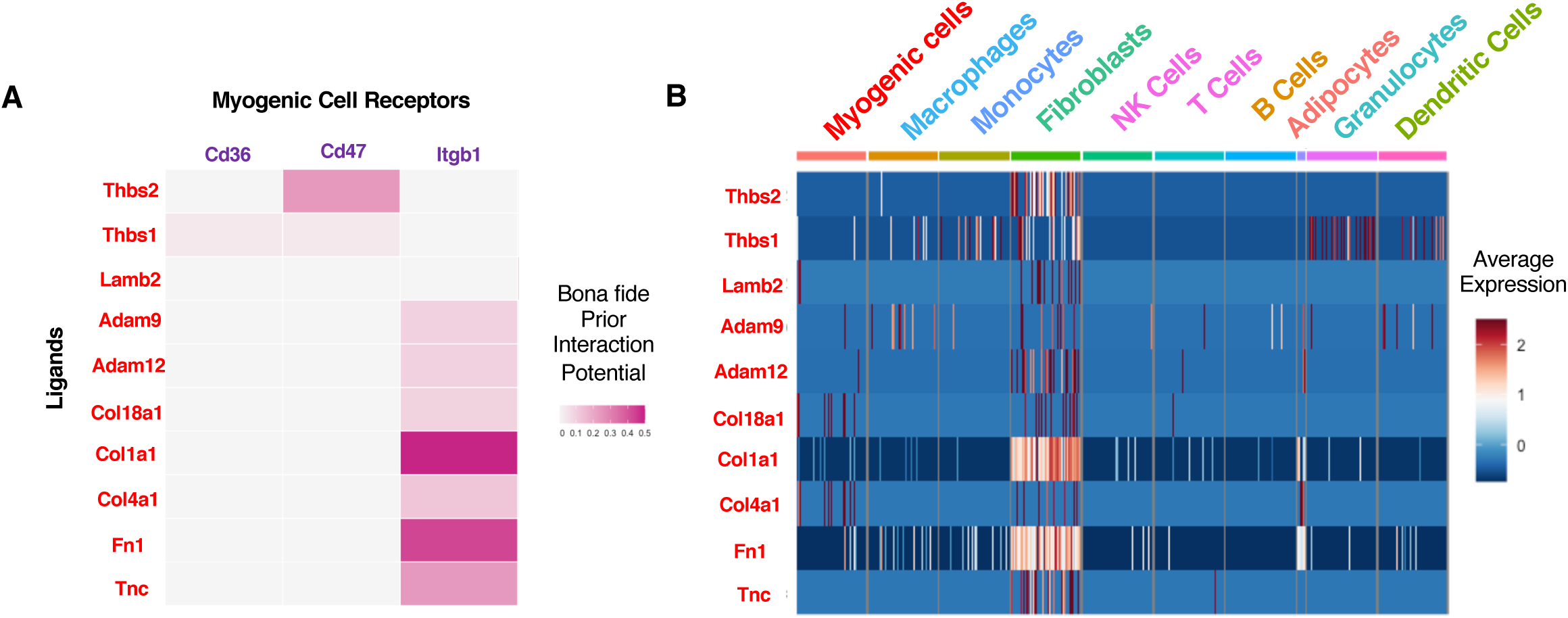
Related to Figure 4. (A) A heatmap depicting the interaction potential of bona fide (present in a validated curated ligand-receptor database), top-ranked NicheNet genes encoding ligands with receptors on myogenic cells. (B) Expression levels for top-ranking bone fide genes encoding ligands in mononuclear cells from regenerating skeletal muscle. The heat map indicates the relative expression level.

**Supplemental Table S1. Differential gene expression analysis of self-renewing myogenic cluster** (Tab 1) Differential gene expression analysis for myogenic Cluster 3, (Tab 2) Differential expression analysis for myogenic cluster 3 versus myogenic cluster 0. The gene with significance, average log fold change, the percentage of cells where the gene is detected in cluster 3, the percent of cells where the gene is detected in the comparison group, and adjusted p-value are provided.

**Supplemental Table S2. Reactome pathway analysis performed on differentially expressed genes in Cluster3.** The pathways, their fold enrichment, the raw p-value, and the false discovery rate are provided in the table. Highlighted pathways indicate pathways that include candidate genes that may regulate MuSC self-renewal.

**Supplemental Table S3. Skeletal muscle ligands and ligand-receptor interaction pairs identified during days 4-7 of muscle regeneration.** (Tab 1) All NicheNet imputed differentially expressed ligands in the regenerating muscle (Tab 2) NicheNet identified ligand with cognate receptor pair inferred from Ramilowski et al, 2015.

**Supplemental Videos 1-9**

Representative 3D renderings of protein immunoreactivity for ParD3 (red) as a marker of asymmetric division, phosphorylated histone 3 (white) as a marker of dividing cells, and self-renewal candidate proteins (green) as well as DAPI dye (blue) to mark nuclei in MuSCs on isolated myofibers. SV_1 Wisp2, SV_2 Secondary alone control, SV_3 Ncoa7, SV_4 mgp, SV_5 mest, SV_6 Itm2a, SV_7 HO-1, SV_8 Gabarap symmetric division, SV_9 Gabarap

